# OmicsOne: Associate Omics Data with Phenotypes in One-Click

**DOI:** 10.1101/756544

**Authors:** Yingwei Hu, Minghui Ao, Hui Zhang

## Abstract

The rapid advancements of high-throughput “omics” technologies have brought huge amount of data to process during and after experiments. Multi-omic analysis facilitates a deeper interrogation of a dataset, and discovery of interesting genes, proteins, lipids, glycans, or metabolites, or pathways related to the corresponding phenotypes in a study. Many individual software tools have been developed to analyze and visualize the data. However, integrating multiple omics data analysis strategies and approaches in a single data processing pipeline is still a challenge task. OmicsOne is a software developed in R, Python and Jupyter Notebook that can achieve statistical analysis, machine learning, and data visualization on multi-‘omics’ data by taking the advantages of integrating the useful tools from individual software packages. OmicsOne can simplify “omics” data analysis, and delineate molecules, or pathways associated to interested phenotypes.

## Introduction

The advancements of high-throughput “omics” technologies, such as genomics, epigenomics, transcriptomics, proteomics, protein modifications, glycomics, lipidomics, and metabolomics, have produced incredible volume of data^1–6^. It is predicted that the trend of generating large datasets will continue as novel technologies are developed and current approaches advance. In this era of omics data explosion, a simple, efficient bioinformatic analysis tool is critical to work with the large amount of data.

Individual tools, such as GiaPronto^7^, PANDA-view^8^, ImageGP (http://www.ehbio.com/ImageGP), offer select options for data analysis and visualization. However, integrating multiple data analysis strategies and approaches in a single data processing pipeline facilitates a deeper interrogation of a dataset, and discovery of interesting molecules or pathways related to the corresponding phenotypes in a study.

Here, we present the tool OmicsOne, a software developed in R, Python and Jupyter Notebook that can preprocess and analyze the multi-“omics” data. After user selected parameters, the results could be reported and visualized in a simple, one-click format. The pipeline includes statistical analysis, machine learning, and data visualization components based on several predefined templates integrated from previous publications^4,6^. OmicsOne can simplify “omics” data analysis, and delineate genes and pathways associated to phenotypes. OmicsOne is hosted on GitHub (https://github.com/huizhanglab-jhu/OmicsOne) and can be downloaded to run locally or on cloud directly via mybinder.org.

## Methods

OmicsOne is developed for intra-omics studies to discover molecular changes and pathways associated with phenotypes. The OmicsOne platform is composed of several essential data processing modules, including preprocessing, profiling, quality control, differential expression analysis, dimensionality reduction, clustering, and visualized reporting (Table 1). The phenotype associations are investigated via the identification of molecules and pathways enriched in phenotypical classifications and statistical calculation of the significant ones. In the following sections, OmicsOne was applied on the public proteomic data set of ovarian cancer to demonstrate the functions^6^.

**Table 1.**
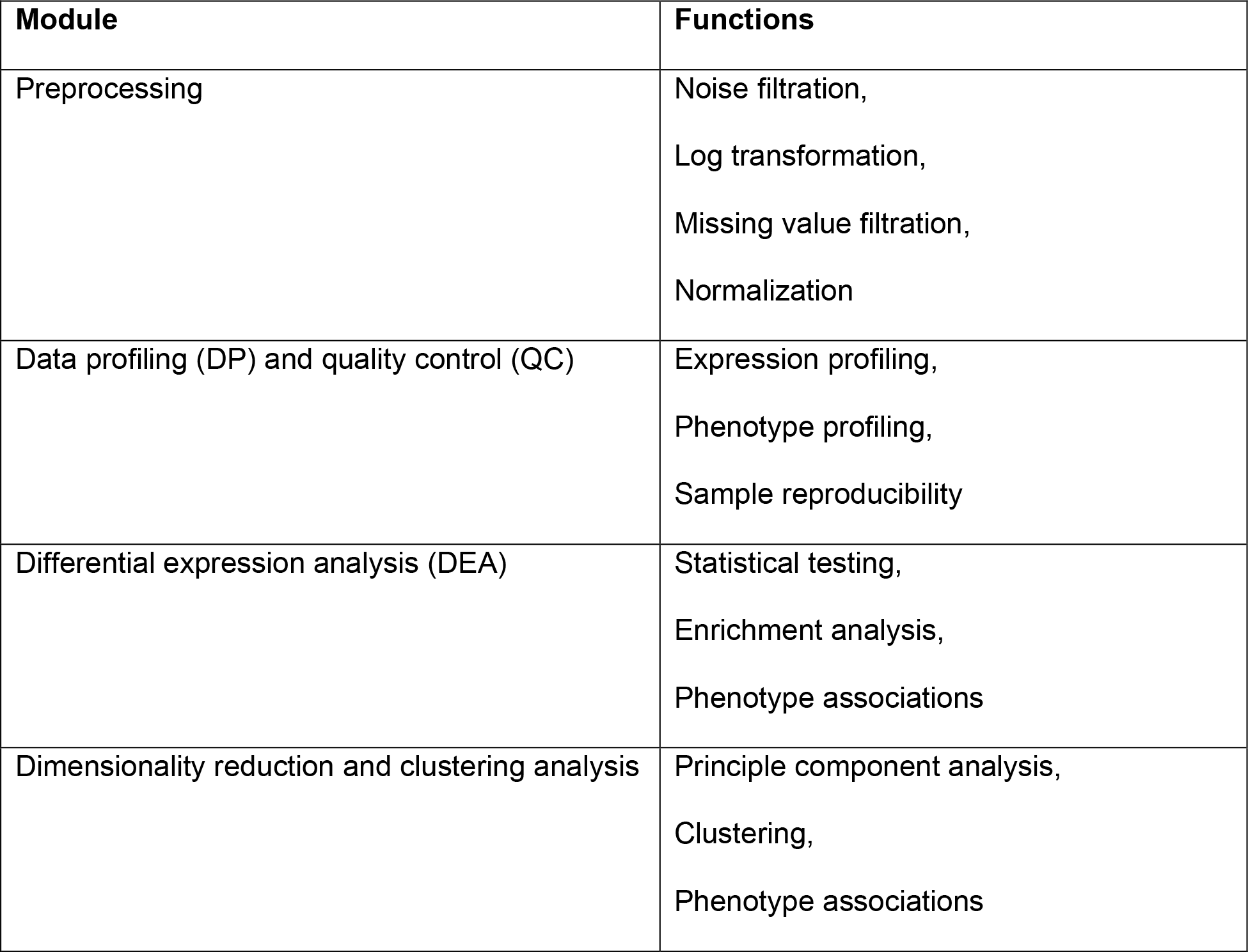
Main modules and functions of OmicsOne

| Module | Functions |
| --- | --- |
| Preprocessing | Noise filtration,<br>Log transformation,<br>Missing value filtration,<br>Normalization |
| Data profiling (DP) and quality control (QC) | Expression profiling,<br>Phenotype profiling,<br>Sample reproducibility |
| Differential expression analysis (DEA) | Statistical testing,<br>Enrichment analysis,<br>Phenotype associations |
| Dimensionality reduction and clustering analysis | Principle component analysis,<br>Clustering,<br>Phenotype associations |

### Input data and preprocess

OmicsOne requires a table of phenotype information and a table of expression matrix of quantitative “omics” data (.csv or .xlsx) both in ‘wide format’. In the phenotype information table, each row represents a sample, while each column represents a phenotype. In the expression matrix, each column represents a sample, while each row represents a feature value, such as gene, protein, peptide, or post-translational modification (PTM) expression level.

The expression matrix will be preprocessed before further analysis if necessary. OmicsOne provides common preprocessing functions, including log-transformation, noise filter, missing value filtration, and normalization.

### Data profiling and quality control

Understanding the data is always the first and critical step for all the following analysis. OmicsOne supports a series of statistical functions (e.g. sum, average, median, standard deviation) to profile samples, features (e.g. proteins), and phenotype information (Figure 1), and visualize the results in various figure types, including bar chart (Figure 1A), heatmap (Figure 1B), and etc. It is also important to evaluate the quality of the data to secure starting analysis from a confident data source. The software uses the calculation of the correlation values of technical/biological replicates (Figure 1C) or coefficient of variance (CV) of selected quality control samples (Figure 1D) to evaluate the reproducibility of measured gene or protein level expression. OmicsOne also offers a correlation assessment to evaluate the concordance of the genomic, transcriptomics, or proteomic expression across multiple samples. In addition, samples or features considered as outliers relative to the larger dataset can be highlighted for additional consideration for inclusion or follow-up analyses.

**Figure 1.**
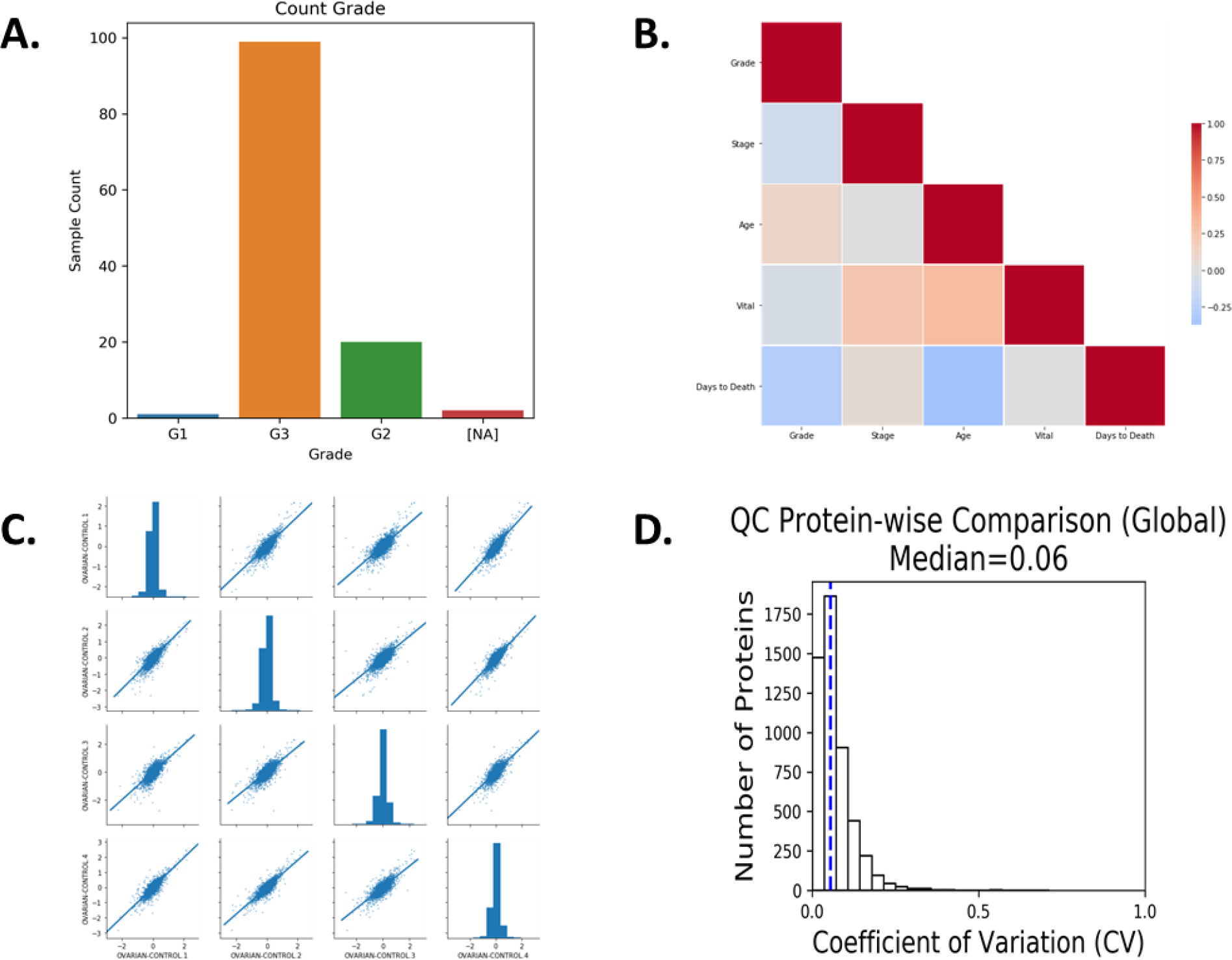
Sample figures of data profiling and quality control module of OmicsOne. **A.** A histogram of grades of all patients. **B.** A heatmap of correlation values of phenotypes to help identify the relationship of known phenotype. **C.** A series of scatter plots with trend lines to demonstrate the reproducibility of quality control samples **D.** A histogram of coefficient of variant (CV) values of protein expression in quality control samples.

### Differential expression analysis (DEA)

Delineating altered expression profiles of genes, proteins, and/or PTMs offer the greatest insight into aberrant biology in comparative studies (i.e. tumor vs. normal, aggressive vs non-aggressive, treated and untreated). OmicsOne can identify the significant, differentially expressed genes or proteins, leveraging multiple statistical tests (t-test, wilcoxon and etc., Figure 2A), followed by subsequent enrichment analysis (GSEApy^9–11^or WebGestaltR^12–15^) to discover up-/down-regulated pathways (Figure 2B). OmicsOne will further calculate the association between the discovered pathways and phenotypes to obtain a comprehensive understanding of the roles of pathways in different conditions (Figure 2C).

**Figure 2.**
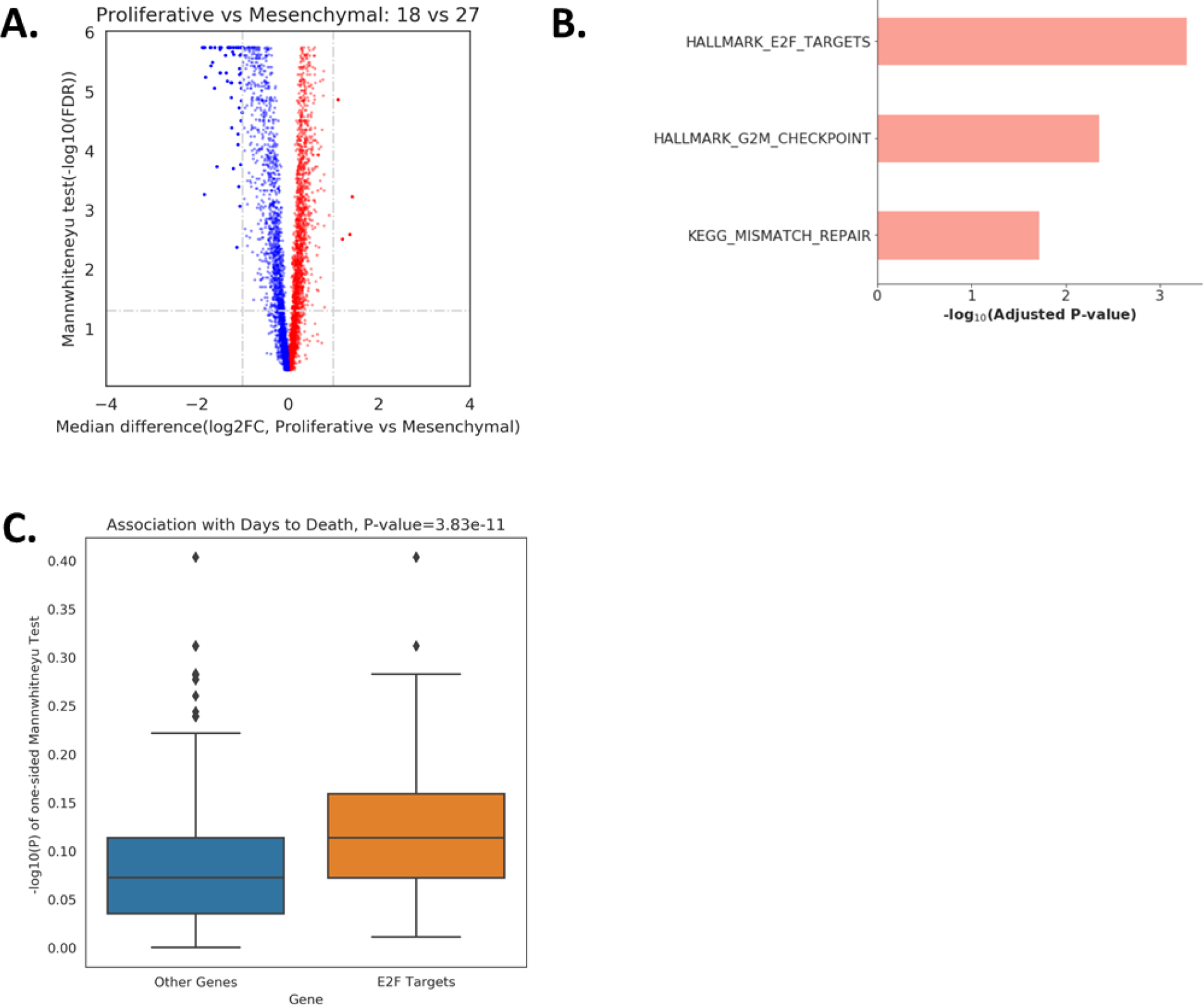
Differential expression analysis module of OmicsOne. **A.** A volcano plot of two ovarian cancer subtypes (proliferative and mesenchymal). The proteins with differential expression were selected as fold change >= 2 and adjusted P values (FDR) < 0.05 **B.** A bar chart of the pathways over-represented in proliferative than mesenchymal, which were enriched from proteins with expression’s fold change >= 1.3 by GSEApy. **C.** A boxplot to show the significant associations of E2F_TARGETS with survival.

### Dimensionality reduction and clustering analysis

Unsupervised methods are useful approaches to classify samples based on the most distinguishable features without prior knowledge; often identifying the most prominent factors driving different phenotypes. Principal component analysis (PCA) and Hierarchical clustering (HC) are two wide-spread methods, supported by Python package: scikit-learn^16^, to cluster samples and identify the signature gene groups associated with the corresponding sample clusters (Figure 3A and 3B). OmicsOne also implemented a customized interface for executing the R packages: CancerSubtypes^17^ and WebGestaltR to evaluate clustering stability and automate gene clustering interpretation. In addition, the significance of identified gene clusters respective to each phenotype or sample cluster can be calculated. Finally, OmicsOne further calculates p-values of associations between the phenotypes and enriched pathways to aid with data application towards biological interpretation.

**Figure 3.**
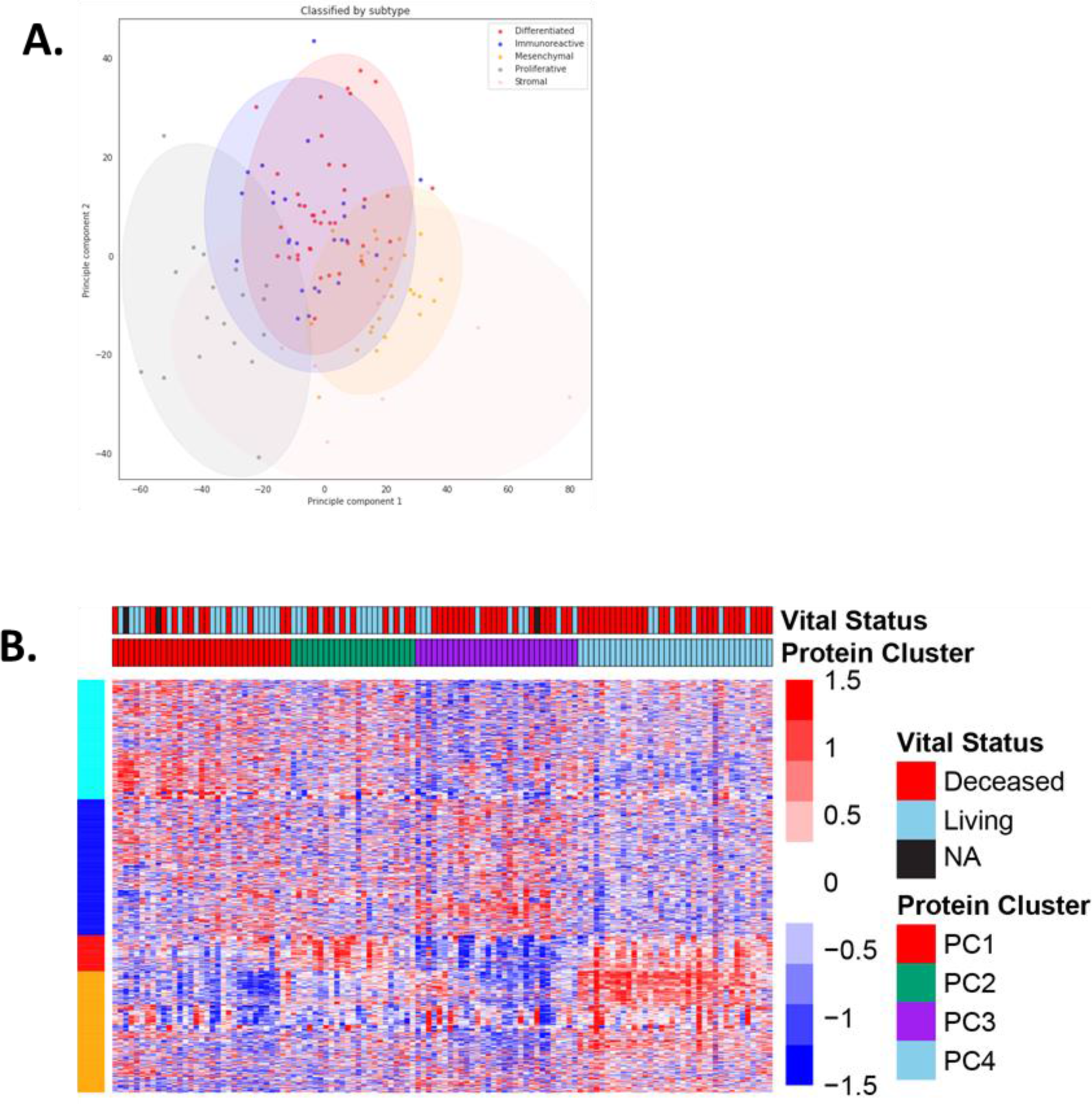
Differential expression analysis module of OmicsOne. **A.** A PCA plot of all 119 samples from Johns Hopkins University (JHU) marked with subtype names and 95% confidence intervals. **B.** A clustering map of all JHU samples with vital status and clustering information analyzed by CancerSubtypes.

### Data visualization and report

All the analysis completed by OmicsOne report intermediate and finalized results in tables (.csv), as well as the corresponding figures. Figures are generated using matplotlib (https://matplotlib.org/) and seaborn (https://seaborn.pydata.org/) packages of Python. OmicsOne defines highly customizable template functions to generate common plotting functions (e.g. bar chart, heatmap, volcano plot, histogram, etc.) to support various approaches of data visualization.

## Conclusion

In summary, OmicsOne is an efficient tool to associate the molecular changes with phenotypes. It was originally designed for isobarically labeled quantitative proteomics data (e.g. tandem-mass-tag (TMT) but can find applications in label-free quantitation and Data Independent Acquisition (DIA) datasets, as well as other “omics” data with proper preprocessing. The software uses predefined templates to build a robust, working pipeline for standard association analyses. Overall, OmicsOne will offer end users a simple, bioinformatic pipeline to identify genes, protein, PTMs, and pathways of interest to understand aberrant biological processes.

## Acknowledgement

This work was supported by the National Cancer Institute, the Clinical Proteomic Tumor Analysis Consortium (CPTAC, Grant U24CA210985) and the Early Detection Research Network (EDRN, U01CA152813). The authors would like to thank Dr. David J. Clark for his assistance with reading the manuscript and helpful discussions

